# Molecular Cross-Validation for Single-Cell RNA-seq

**DOI:** 10.1101/786269

**Authors:** Joshua Batson, Loïc Royer, James Webber

## Abstract

Single-cell RNA sequencing enables researchers to study the gene expression of individual cells. However, in high-throughput methods the portrait of each individual cell is noisy, representing thousands of the hundreds of thousands of mRNA molecules originally present. While many methods for denoising single-cell data have been proposed, a principled procedure for selecting and calibrating the best method for a given dataset has been lacking. We present “molecular cross-validation,” a statistically principled and data-driven approach for estimating the accuracy of any denoising method without the need for ground-truth. We validate this approach for three denoising methods—principal component analysis, network diffusion, and a deep autoencoder—on a dataset of deeply-sequenced neurons. We show that molecular cross-validation correctly selects the optimal parameters for each method and identifies the best method for the dataset.

---

High-throughput single-cell RNA sequencing (scRNA-seq) has become an essential tool to study cellular diversity and dynamics, enabling researchers to discover novel cell types^1^, map whole-organism transcriptomic atlases^2^, describe features of fate determination in development^3–6^, and uncover transcriptional responses to stimuli^7^.

The data from scRNA-seq experiments are distinguished by their sparsity: in cell types where mRNA from ten thousand genes are observed in bulk data, mRNA from only a few thousand of those genes will be detected in each cell. While some of that variability may reflect biological phenomena such as the existence of sub-populations of cell types or transcriptional bursting, much of it is an inevitable consequence of the numbers involved. In high-throughput methods, many cells are sequenced to a depth of only a few thousand unique mRNA molecules^8–10^. Since a typical mammalian cell contains hundreds of thousands of mRNA molecules^11^, many genes present at low levels will not be detected simply by chance^12, 13^. Significant processing and analysis is required to extract biological meaning from such sparse and noisy data.

Many computational methods have been proposed to reduce the noise in single-cell RNA-seq data. The most common approach, principal component analysis (PCA), approximates the gene expression matrix as a product of low-rank matrices, and is included as a preprocessing step in popular software packages for scRNA-seq analysis^14–16^. Deep autoencoders provide a flexible extension to PCA, allowing for hierarchical features, nonlinear effects, and loss functions tailored to count data^17, 18^. In contrast, diffusion-based methods perform local smoothing, where the expression of each cell is averaged with those of the most similar cells from the same experiment^19^. Applying one of these denoising methods can make subsequent analyses simpler: instead of having to design a clustering, trajectory, or pathway method tailored end-to-end to undersampled scRNA-seq data, one may apply a more straightforward method to the denoised data.

Each of these denoising methods has parameters that control the trade-off between removing noise and blurring the biological signal. For PCA, the key parameter is the number of principal components. As more components are used, more of the true variability between cells is retained, but so is more of the noise. For deep autoencoders, the key parameter is the width of the bottleneck layer. When the bottleneck layer is wide, the autoencoder will let noise through, and when the bottleneck layer is narrow, the autoencoder will discard signal^∗^. For diffusion-based methods, the key parameter is how much to diffuse: too little and noise remains, too much and the true heterogeneity of the cells is obscured. Choosing these parameters well can improve all downstream biological analyses.

We propose an approach for estimating the relative accuracy of any single-cell denoising method, inspired by self-supervised approaches to image denoising^20, 21^. We randomly apportion the counts from each cell into two groups, each simulating a shallower but statistically independent measurement of the original cell (Fig. 1a). The accuracy of a denoising model can be evaluated by fitting it on one of the groups (training) and comparing the denoised output to the other group (validation). We prove that this “molecular cross-validation” (MCV) loss approximates, up to a constant, the ground-truth loss, defined as a hypothetical comparison to the full mRNA content of the original cell (Fig. 1b, Methods). Just as ordinary cross-validation may be used to select good parameters for a predictive model on a given dataset, molecular cross-validation may be used to find good parameters for a denoising model.

**Figure 1:**
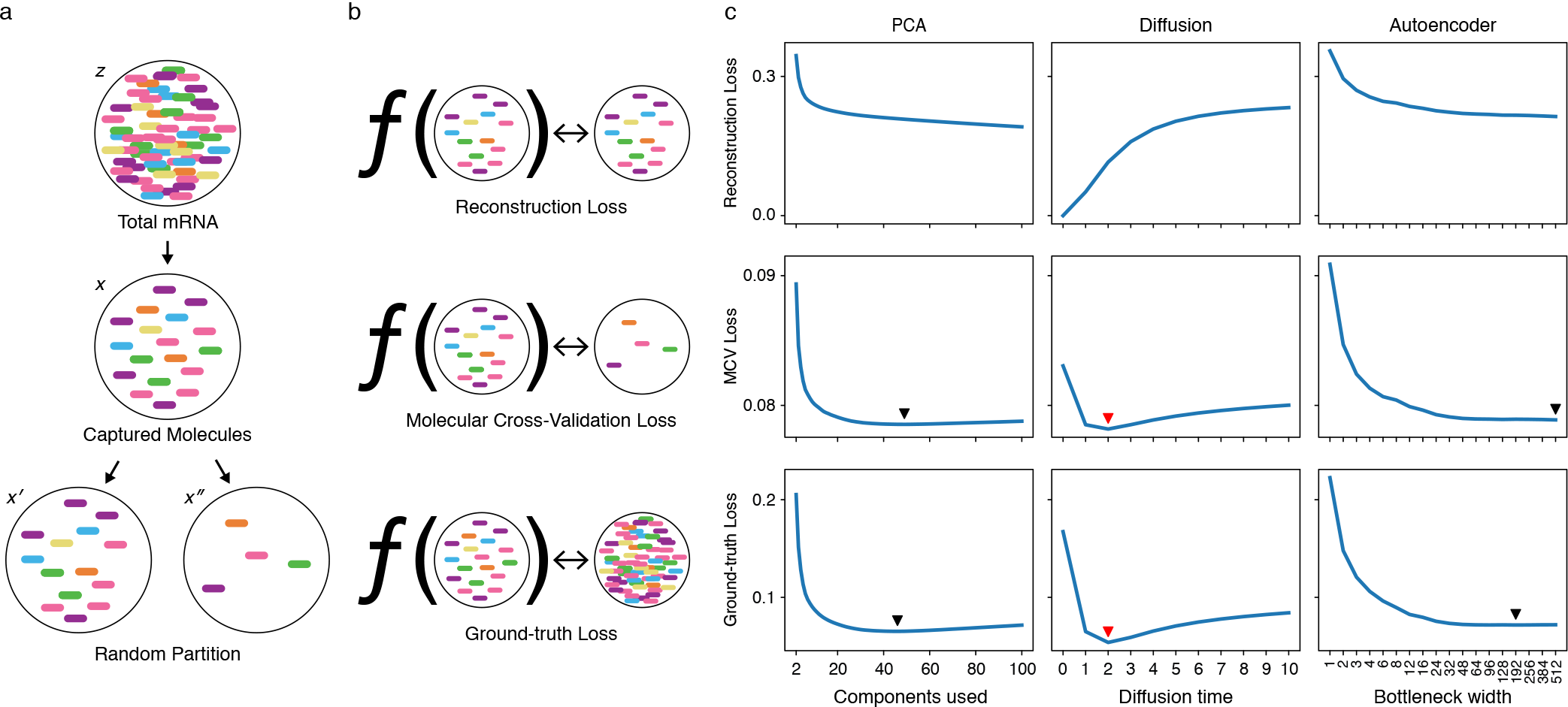
Molecular Cross-Validation loss calibrates denoising methods for single-cell RNA-seq. **(a)** A random partition of the molecules captured from a cell (*x*) simulates two independent samples (*x*′, *x*″) of the cell’s total mRNA (*z*). **(b)** The performance of a denoiser *f* is evaluated in three ways: by comparing *f* (*X*′) to *X*′ (reconstruction loss) and to *X*″ (MCV loss), and by comparing *f* (*x*) to *z* (ground-truth loss). **(c)** Performance of three denoising methods across a range of model parameters. Arrows denote the minima of the MCV and ground-truth loss curves; red arrows denote the global minima among all three methods. Curves shown are for the *Neuron* dataset of 4581 deeply-sequenced cells.

We validate this approach on two datasets for which we have a form of ground-truth. The first (*Neuron*) is a set of 4581 deeply-sequenced neurons from a large dataset of 1.3 million neurons^22^. We selected all cells with at least 20,000 UMIs and subsampled them to 3000 molecules to simulate a typical shallow-depth experiment. We split molecules into training and validation sets using a 90/10 ratio, and compute the average loss over ten splits. The original counts serve as a proxy for ground-truth gene expression. In Figure 1c, we show how three key metrics vary as we sweep the parameters of three different denoising models on the *Neuron* dataset. The second example is a simulated dataset of 4096 cells where the molecules detected are drawn at random from a known distribution. The corresponding plots for the simulated dataset are in Supplementary Fig. 1.

The first metric shown, the reconstruction loss, measures the difference between the training data and the denoised training data (Fig. 1c, top row). This is the objective function explicitly minimized by PCA and the deep autoencoder, and it strictly decreases as the number of principal components or bottleneck width increases. In diffusion-based methods, the reconstruction loss strictly increases as a function of diffusion time, where *t* = 0 leaves the input data unchanged and *t* >> 0 corresponds to replacing each cell with the bulk average. Because the reconstruction loss vanishes when no denoising is done, it is a poor measure of the quality of a denoiser.

The second metric is the molecular cross-validation loss, which measures the difference between the validation data and the denoised training data (Fig. 1c, middle row). The third metric is the ground-truth loss (Fig. 1c, bottom row), which measures the difference between the denoised data and the true gene expression of the original cell. The best parameter values for a particular model are those which minimize the ground-truth loss.

In accordance with the theory, the MCV loss and the ground-truth loss curves have almost identical shapes^†^. In particular, their minima occur at nearly the same parameter values. That means one can select the optimal parameters for each denoising algorithm by finding the minimizer of the molecular cross-validation loss (Fig. 1c check marks). One can also use the MCV to decide between algorithms. For the *Neuron* data, the MCV loss is minimized by diffusion with *t* = 2; note that this is also the minimizer of the ground-truth loss.

The losses above are based on the mean-square error, a generic loss function for any numerical data. For count-based data such as scRNA-seq, one may also use a count-based loss such as Poisson. The MCV procedure extends naturally to the Poisson loss, and in the case of the *Neuron* dataset, selects the same optimal method and parameter value (Methods, Supplementary Fig. 2).

The qualitative effect of choosing the right parameter values for a denoiser is illustrated in Figure 2. We calibrate PCA on a dataset (*Myeloid*) of 2417 myeloid bone marrow cells from Paul, Arkin, & Giladi *et al.*. In Fig. 2a we show clustered heatmaps for 33 genes of interest in this dataset. When only the first three components are used (far right panel), the gene expression is forced into a block structure, artificially removing heterogeneity that was present in the original sample. Conversely, when too many components are used, sampling noise is retained and some subtle relationships are lost. In Fig. 2b we show that the qualitative relationship between the expression of *Gata1* and *Apoe* depends on the amount of smoothing. By selecting an optimal number of principal components with MCV, it is possible to see separation between the *Gata1*-low and -high populations. When too many components are used, the noise overpowers this signal and the presence of *Apoe* in a *Gata1*-intermediate population is difficult to discern. When too few components are used, only a broad and perhaps spurious correlation is visible.

**Figure 2:**
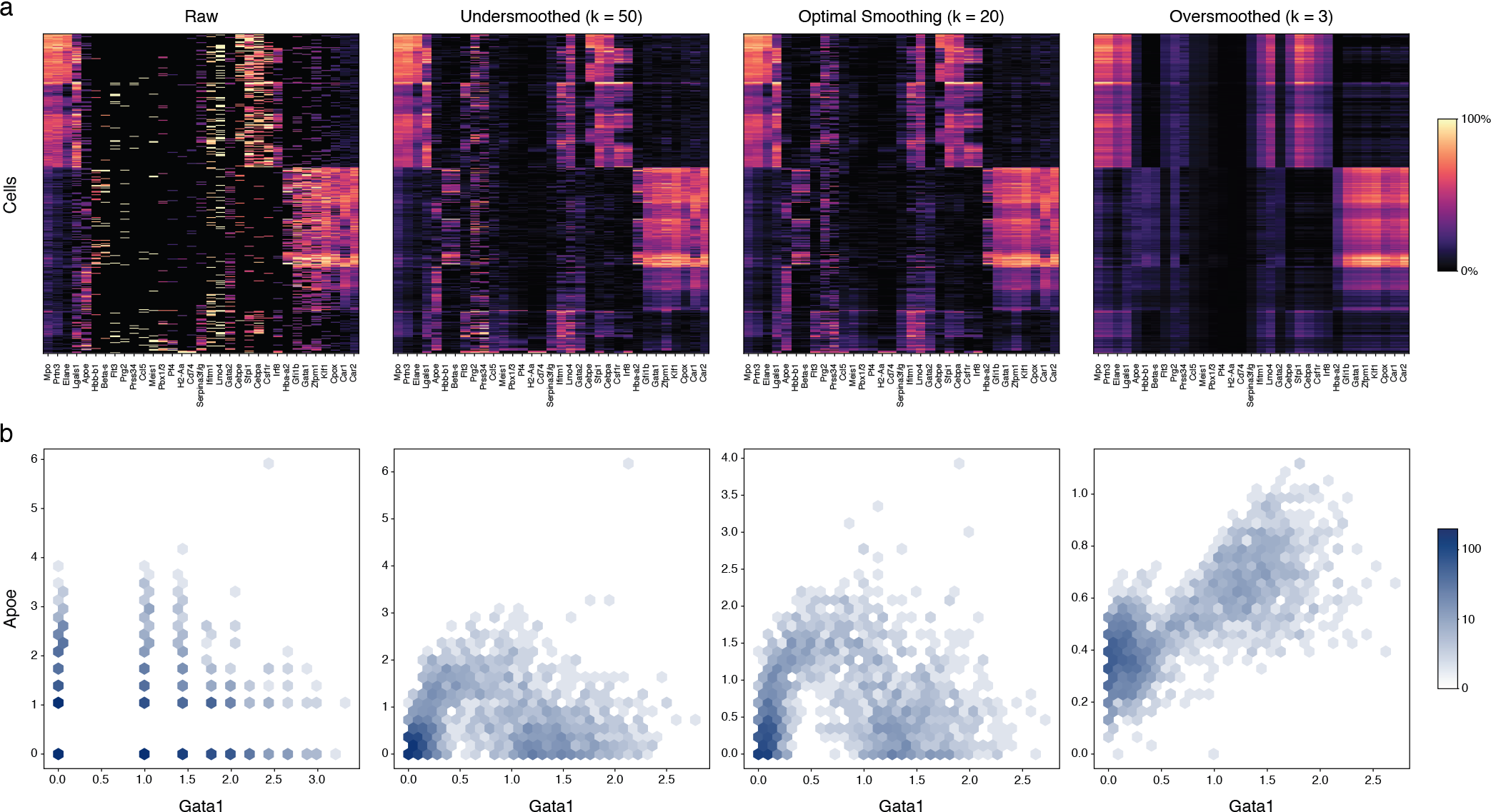
The effect of denoising bone marrow data from the *Myeloid* dataset using PCA with different numbers of principal components. **(a)** Heatmaps for 33 genes of interest. Expression values have been scaled per-gene and re-ordered based on the optimally-smoothed expression matrix. **(b)** Hexbin plots showing the relationship between *Apoe* and *Gata1* in the data at each level of smoothing.

PCA is a simple method for denoising, with one free parameter. More complex methods can have many more parameters, and the MCV loss may be used to simultaneously calibrate all of them. We demonstrate this for MAGIC, which combines PCA with graph diffusion^19^, on a dataset of cells undergoing an epithelial-to-mesenchymal transition. We focus on three genes which provide a portrait of the transition: an epithelial marker, *CDH1*, a mesenchymal marker, *VIM*, and a transcription factor, *ZEB1*. We perform a grid search to find the values of three parameters of MAGIC which are best for these data: the number of principal components, the number of neighbors used to build the graph, and the diffusion time. We find that the optimal parameter values (20 PCs, 4 neighbors, and 1 diffusion step) differ appreciably from the default parameters (100 PCs, 10 neighbors, and 7 diffusion steps). In Figure 3, we show the qualitative consequences of that difference. While no relationship between the three genes is discernible in the raw data, the version denoised with default parameters shows a strikingly smooth transition from a *CDH1*-high *VIM*-low state to a *CDH1*-low *VIM*-high state, with *ZEB1* turning on somewhere in the middle. When denoised with optimal parameters, however, the data reveal a more heterogeneous version of the same general trend. At low values of *CDH1*, there is a wide range of *VIM* and *ZEB1* expression, perhaps representing the natural variation exhibited by cells along this trajectory. This demonstrates the danger of evaluating denoising methods by agreement with expected patterns. The patterns of gene expression learned from bulk studies will appear more clearly as data is oversmoothed.

**Figure 3:**
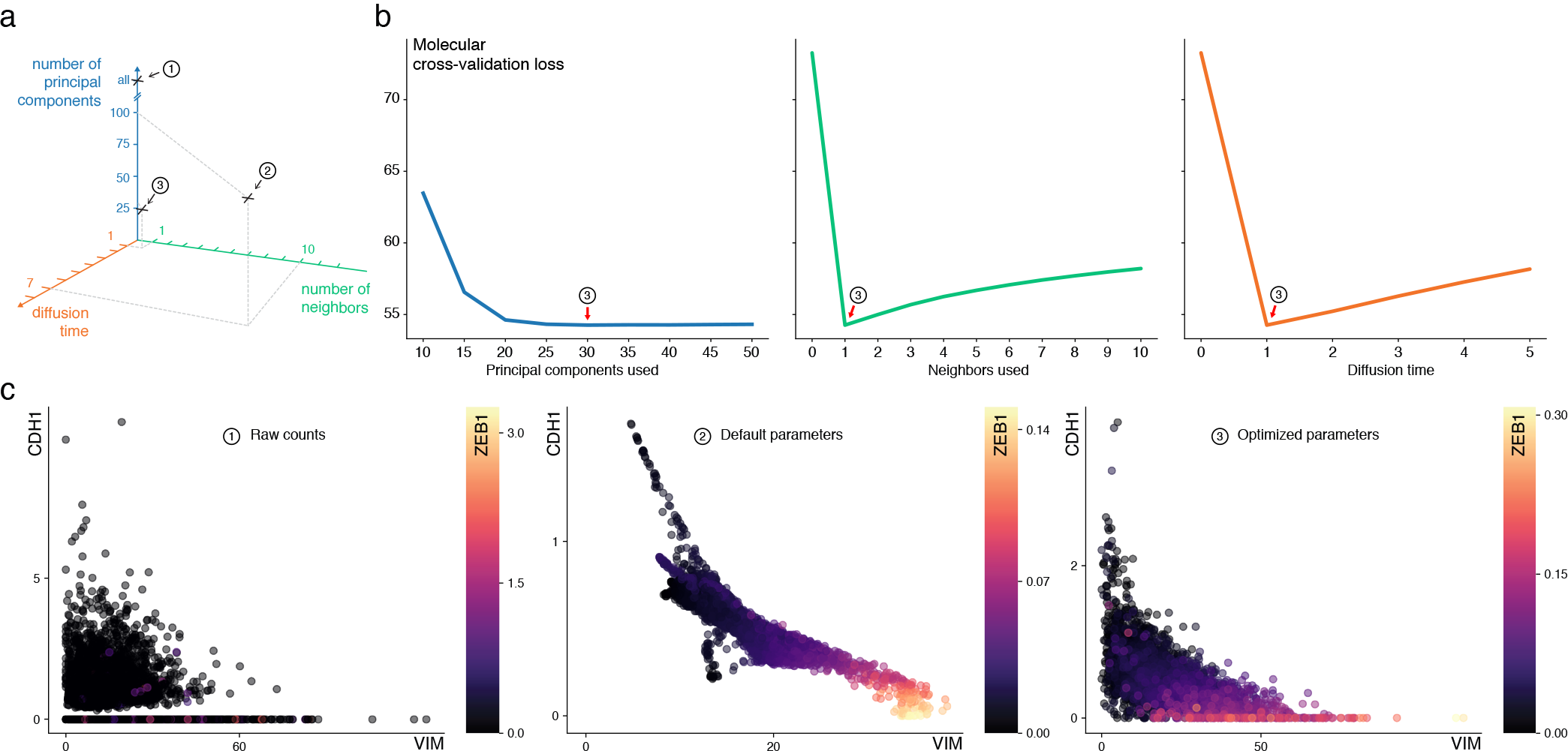
MAGIC denoising algorithm with parameters optimized using the MCV loss shows a heterogeneous portrait of the epithelial-mesenchymal transition. **(a)** Parameter space for MAGIC algorithm. Each point represents a setting for diffusion time, the number of graph neighbors, and the number of principal components. **(b)** Effect on MCV loss of changing each parameter away from optimal value (red arrows). **(c)** The relationship between an epithelial marker, *CDH1*, a mesenchymal marker, *VIM*, and a transcription factor, *ZEB1*, is indiscernible in the raw data, overly smooth when denoised with default parameters, and present but heterogeneous when denoised with optimal parameters.

## Discussion

In this work we demonstrate molecular cross-validation, an approach for evaluating any method for denoising single-cell RNA-seq data. As more tools for scRNA-seq analysis become available, there is an increasing burden on researchers to run, tune, and evaluate the performance of different methods on their specific data. This process is time-consuming and prone to bias, as it is tempting to select the method giving the best concordance with prior biological knowledge. In contrast, molecular cross-validation provides an unbiased way to both calibrate a given denoising method and to compare its performance to other methods. This allows researchers to take advantage of novel methods when they offer better performance on their data.

The key feature of molecular cross-validation is that it directly estimates the quantity of interest: the similarity of the denoised data to the full set of mRNA present in the original cell. This avoids the pitfalls of existing approaches to calibration. Some approaches are specific to certain models: bi-cross-validation^23^ and eigenvalue-localization^24^ only apply to matrix factorization models like PCA. Information-theoretic approaches like the Akaike information criterion fail to extend to nonparametric methods like diffusion or overparameterized models like deep neural networks^25^. Finally, metrics that measure the concordance of downstream results with prior expectations, such as silhouette width, do not directly reward accuracy^26^. For example, replacing the expression of each cell with the average expression of its cluster will increase the silhouette width while removing all within-group heterogeneity. The MCV loss, in contrast, makes no assumptions about the structure of the model or the dataset.

The accuracy of the estimate of the ground-truth loss provided by a single round of MCV depends on the choice of train-validation split. As more molecules are used for training, the optimal parameters for the training data will approach the optimal parameters for the full data. However, as fewer molecules are used for validation, the MCV loss will be less stable over different random splits. To resolve this bias-variance tradeoff, we compute the average of the MCV loss over many random splits (here, 10), using most of the molecules for training each time (here, 90%). There may be computationally efficient ways to adaptively select the training-validation split and number of replicates; this is an interesting topic for future work.

We have shown how to calibrate the parameters of *deterministic* single-cell denoising methods, and how to select the best method for a given dataset. Probabilistic methods for modeling single-cell data, such as scVI and SAVER^18, 27^ also have parameters that require tuning, and calibrating and comparing such methods is an interesting direction for future work. It is also possible that more complex methods with many more parameters could be developed, using the MCV loss to fit those parameters in a principled way. One might even train a model to directly minimize the MCV loss, as in recent self-supervised deep learning models for image denoising^21, 28^.

## Acknowledgements

We would like to thank Casey Greene, Dana Pe’er, Rahul Satija, Jingshu Wang, Andre Wibisono, and Nancy Zhang for valuable discussions, and `your name here` for valuable comments on the manuscript. Funding for this work was provided by the Chan Zuckerberg Biohub.

## Correspondence

Correspondence and requests for materials should be addressed to.

## Code

Available on GitHub at www.github.com/czbiohub/molecular-cross-validation.

## Methods

We begin by describing a simple statistical model of the capture and sequencing process. Consider a collection of cells *c*_1_, …, *c*_*n*_, each containing a set of mRNA molecules Ω_1_, …, Ω_*n*_. The output of a single-cell RNA-seq experiment using unique molecular identifiers (UMIs) will be sequences from a random subset of the molecules from each cell. We assume that the molecules detected from cell *i* are drawn uniformly at random from Ω_*i*_, with each molecule having a probability *p*_*i*_ of being detected. (The capture efficiencies *p*_*i*_ may differ between cells.) By aligning the sequences of detected molecules to a genome, a vector *x*_*i*_ of counts for each gene is produced.^‡^ We write *X* for the full cell-by-gene matrix, where *X*_*ij*_ is the count of molecules from cell *i* mapping to gene *j*. We also consider a hypothetical matrix *X*^*deep*^ which would have been produced if all of the molecules from each cell were detected. If we imagine rerunning the random capture process on the same set of cells, then *X* becomes a random matrix with entries 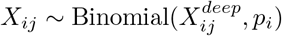.

It is common in the literature to use a Negative Binomial distribution to model the variability in gene counts between cells^17, 18^. This represents three kinds of variability: biological differences between cells, variability in library size, and sampling. Here, we are taking the mRNA content of each cell and the fraction of molecules captured as fixed. The only remaining variability is in *which* molecules are sampled, yielding the Binomial distribution above. Note that for sequencing methods without UMIs, counts do not represent independently captured molecules, violating the assumptions required for molecular cross-validation.

We view a denoising algorithm as a function *f* which takes in the entire matrix of observed counts *X* and produces an estimate of *X*^*deep*^. We would like to select a function *f* for which the loss 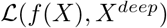 is small, for an appropriate loss function 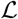. Molecular cross-validation (MCV) is a procedure for estimating that loss (up to a constant) from *X* alone. Before describing MCV, we first recall some properties of ordinary cross-validation (CV) and the difficulties of applying CV to the task of denoising.

### Cross-Validation

In an ordinary prediction problem, one fits a model *g* to a training set 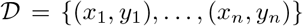. The inferential task is to estimate the accuracy that the resulting predictor 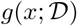 would have on new data points. Usually, one is considering some family of models *g*_*m*_ with a hyperparameter *m*, and is looking to select the model which will generalize the best.

Note that the error of the model on the training data itself,

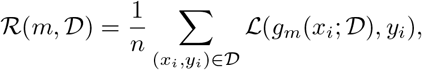

is a poor prediction of generalization, as more complex models will always fit their training data better.

In cross-validation, one repeatedly splits the training dataset into complementary pieces 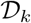 and 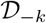, each time using the piece 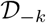 to train the model and the smaller piece 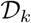 to evaluate it. [In traditional *K*-fold cross-validation, each training data point appears in exactly one 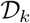. Similar estimates may be obtained by choosing each 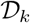 to be a random subset of 1/*K* of the training data.] The estimate of the generalization error is

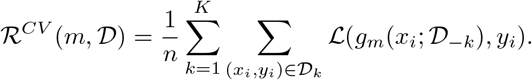

This is an imperfect estimate for the generalization error of 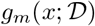, since each time the model is fit it has access to only a subset of the training data. Perhaps a more expressive model could be fit if all of the data were used, but in the limit of large training sets, this gap disappears.

Difficulties arise when cross-validation is used on an unsupervised task like denoising, where only the noisy data *x* is observed. One may still use CV to estimate the expected reconstruction error 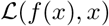 for a model *f* by fitting it on some data points and validating it on held-out data points. However, the expected reconstruction error is not a measure of the quality of the denoiser, as the identity function, which does no denoising, will generate zero loss on both train and validation sets. Since a more complex model will be better able to approximate the identity function, it will have lower loss on the held-out validation sets even when it has over-fit the data. For example, PCA with *m* principal components fits a model of the form *g*_*m*_(*x*) = *UV*^*T*^_*x*_ where *U* and *V* are *m* × *p* matrices. As *m* increases, the capacity of the model goes up, and 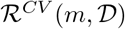 strictly decreases. However, the distance between 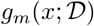 and the ground truth will decrease until an optimal *m* is reached and then increase thereafter (as in Figure 1).

To get a proper estimate of the denoising quality of *f*, we need a way to decouple the noise in the data used to fit *f* from the noise in the data used to evaluate it, so that the identity mapping will no longer be optimal.

### Independence

While we may never have access to the ground truth *X*^*deep*^, we can construct statistically independent samples from it by carefully splitting our measurement *X*.

If we did have two independent random captures of a cell with efficiency *p*′ and *p*″, then their overlap would be a capture of efficiency *p*′*p*″ and their union would be a capture of efficiency

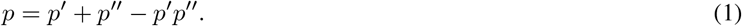

We instead begin with a single capture *S* of efficiency *p* from a set of molecules Ω, and choose probabilities *p*′ and *p*″ satisfying (1). Then we work backwards to generate independent samples summing to the observed sample: we randomly partition the captured molecules into three groups *S*_1_, *S*_2_, *S*_3_ with relative proportions *p*′(1 − *p*″): *p*′*p*″: *p*″(1 − *p*′). The unions *S*′ = *S*_1_ ⋃ *S*_2_ and *S*″ = *S*_2_ ⋃ *S*_3_ form independent draws from Ω with union *S*. The corresponding formulation for the cell-by-gene matrix of observed counts is as follows:

#### Proposition 1.

*Fix a count matrix X*^*deep*^, *capture efficiencies p*_*i*_, *and a validation ratio α*. *Then there exist probabilities* 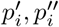 *such that if we draw*

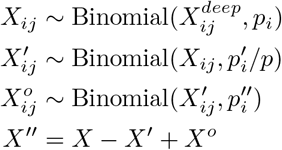

*then X′ and X″ are independent random variables with entries distributed* 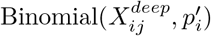 *and* 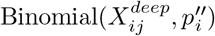, *repestively, and* 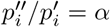.

*Proof*. The conditions 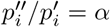 and *p* = *p*′ + *p*″ − *p*′*p*″ produce a quadratic equation for *p*′:

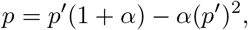

whose solution is

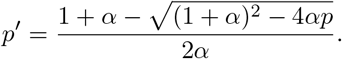

We then set *p*″ = *αp*′. The claim follows from analyzing the conditional probabilities. Take two independent draws *S*′ and *S*″ from a set Ω with probabilities *p*′ and *p*″, where now 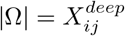, and set *S* = *S*′ ∪ *S*″. Then for a given molecule *m*, we have

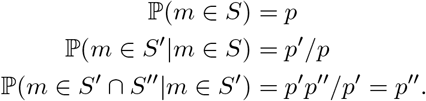

The principal of inclusion-exclusion for the sizes of the sets *S*, *S*′, and *S*″ finishes the proof.

Because the noise in the renditions of each cell in *X*′ and *X*″ is independent while the underlying signal is the same, training a denoising model on one and validating its output using the other gives insight into its accuracy.

In what follows, we suppress indices *i*, *j* where both sides of an equation straightforwardly vectorize.

#### Proposition 2 (MSE Loss).

*Let X* ~ Binomial(*X*^*deep*^, *p*). *Fix a validation ratio α, and let X*′, *X*″ *be random splits of X and let p*′, *p*″ *be probabilities as in Proposition 1. Let f be an arbitrary denoising function. Then*

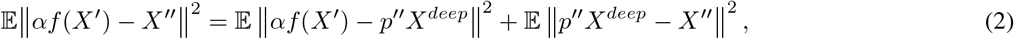

*where the expectations are with respect to the sampling of X from X*^*deep*^ *and X*′, *X*″ *from X*.

*Proof.* We expand the left-hand side as,

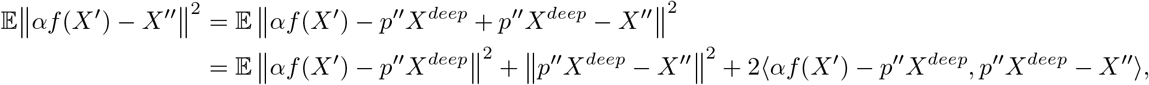

where ⟨*A*, *B*⟩ denotes the entrywise inner product between matrices. By Proposition 1, the process of drawing *X* from *X*^*deep*^ and splitting it into *X*′ and *X*″ is equivalent to drawing *X*′ and *X*″ independently from *X*^*deep*^ and adding them (less overlap) to get *X*. Since *X*′ and *X*″ are independent, we may bring the expectation inside the third term:

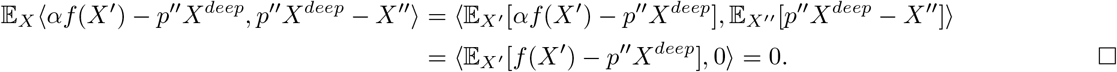

In the formulation above, *f* can be an arbitrarily complex function of the input matrix *X*′. For example, *f* may be “perform PCA on *X*′ and project it onto the first *k* principal components” or “train a deep autoencoder neural network using stochastic gradient descent with a cosine annealed learning rate and weight decay with random seed equal to 42.”

Equation 2 states that the molecular cross-validation loss (left-hand side) is equal to the ground-truth loss (right-hand side, first term) plus a constant independent of *f* (right-hand side, second term). The denoising function minimizing the MCV loss will also be the function minimizing the ground-truth loss.

### Practical Considerations

Note that the ground-truth loss on the right is for the denoiser applied to *X*′. If one chooses parameters for *f* which minimize the MCV, they will be optimal for *X*′ but not necessarily for the full set of molecules *X*. In a typical usage, where 90% of the molecules are used for training (corresponding to *α* = 1/9), *X*′ is close to *X* and it is reasonable to expect that the optimal parameters for denoising *X*′ will be close to those for denoising *X*. In practice, we find this to be the case (e.g. Figure 1 and Supplementary Figure 1.) This is analogous to the situation for ordinary CV, in which optimal hyperparameters for fitting a model on 90% of the data points may be slightly suboptimal for fitting a model on all of the data; a more complex model might take advantage of having more data to fit on. Nevertheless, it is common practice to take the hyperparameters found using CV and use them to fit a model on all the data, and we recommend the same procedure for MCV.

Note that the overlap between the molecules in *X*′ and *X*″ in Proposition 1 is very small when the capture efficiency *p* is low. For a cell with 5000 molecules detected from a population of 500,000 (*p* = 1%) and validation ratio *α* = 1/9, the expected overlap is only 4.5 molecules. **In practice, one may simply partition the molecules in** *X*, setting *X*′ ~ Binomial(*X*, 1/(1+*α*)) and *X*″ = *X* − *X*′. On the other hand, if a significant fraction of the molecules from *X*^*deep*^ are captured, one should use an overlap as in Proposition 1. This is the case for the *Neuron* dataset below, where the deeply sequenced cells used as a proxy for ground-truth contain as few as 20,000 molecules.

### Normalization

To cope with differing capture rates between cells and differing magnitudes of expression for different genes, it is common to normalize gene expression matrices. For example, the rows of the matrix may be normalized to “counts per *N*” for some *N*, and the resulting matrix entries may be log or square-root normalized. Molecular Cross-Validation can also be used to estimate the distance of a denoised normalized matrix to an appropriately normalized ground truth. The appropriate ground truth is the expected value of the normalized matrix, which, since expectations do not commute with nonlinear functions, is not given by naively normalizing a downscaled deep count matrix. Because the nonlinear effects of normalization may be different at different sampling depths, specifically on the training and validation versions of each cell, a rescaling function may be necessary to convert the denoised training matrix to the scale of the validation matrix.

#### Proposition 3 (MSE Loss with Normalization).

*Let X* ~ Binomial(*X*^*deep*^, *p*). *Fix a validation ratio α, and let X*′, *X*″ *be random splits of X and p*′, *p*″ *be probabilities as in Proposition 1. Let f be an arbitrary denoising function, let η be an arbitrary normalization function, and let v*: ℝ → ℝ *be an arbitrary rescaling function. Then*

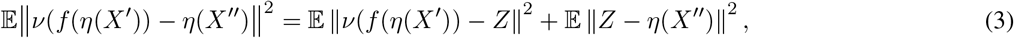

*where the expectations are with respect to the sampling of X from X*^*deep*^ and *X*′, *X*″ *from X. where* 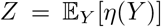 *for Y* ∼ Binomial(*X*^*deep*^, *p*″).

*Proof.* As before, we exploit the independence of *X*′ and *X*″. We have,

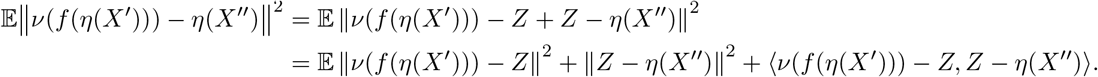

Since *X*′ and *X*″ are independent, we may bring the expectation inside the third term:

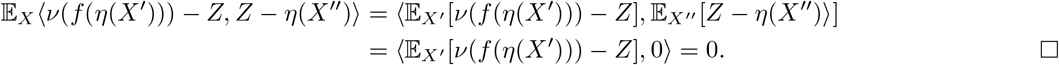

In the case where no normalization is used (i.e., *η* is the identity function), then *Z* = *p*″*X*^*deep*^, the scaling function *v* is multiplication by *p*″/*p*′ = *α* and this reduces to Proposition 2.

For square-root normalization, the situation is more subtle. To see this, consider a cell *i* with 10 molecules of gene *j*, so 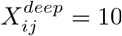. Then the appropriate ground-truth *Z*_*ij*_(*p*″) is a nonlinear function of *p*″: For example, at *p*″ = 1 it is 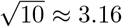, at *p*″ = 0.1 it is 0.79, and at *p*″ = 0.01 it is 0.10. In theory, the appropriate rescaling *v* would depend on *n*, *p*′, and *p*″, but when *p*′ is relatively small, i.e., most molecules are not captured, the binomial distribution can be approximated by a Poisson distribution and the calculation simplifies considerably. If we set

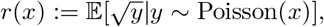

then we have *v*(*x*) = *r*(*αr*^−1^(*x*)). This is the rescaling used to post-process the output of all square-root normalized denoisers when computing the MCV loss.

### Poisson Loss

The Mean-Square Error loss is quite useful, as the corresponding models are easy to fit and its minimizer is the expected value of the target. However, the Poisson loss is a closer match to the generating process of the data. The log-likelihood of observing *k* in a Poisson distribution with mean *μ* is loglik(*μ*, *k*) = *μ* − *k* log *μ*. We use the same notation to describe the total log-likelihood for a vector or matrix of means and a matrix of counts:

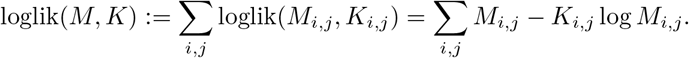

Unlike for the MSE, the Poisson version of the MCV loss has no term for the noise variance: the MCV loss is equal in expectation to the appropriate ground-truth loss.

#### Proposition 4 (Poisson Loss).

*Let X* ~ Binomial(*X*^*deep*^, *p*). *Fix a validation ratio α, and let X*′, *X*″ *be random splits of X and p*′, *p*″ *be probabilities as in Proposition 1. Let f be an arbitrary denoising function. Then*

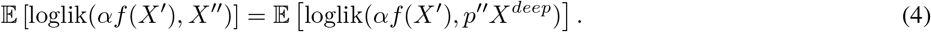

*Proof.* As before, we exploit the independence of *X*′ and *X*″. We have

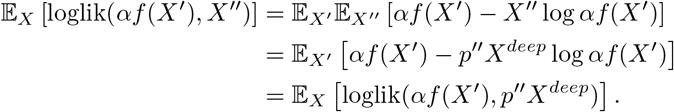

Since *f*(*X*′) is independent of *X*″, and the Poisson loss is linear in the count variable, the expectation with respect to *X*″ can be evaluated inside the loss without affecting the other terms.

More generally, MCV will work for any loss function which is the log-likelihood of an exponential family. This includes the mean-square error, which is the log-likelihood of a Gaussian, and the Poisson loss, as above, but also the Negative Binomial distribution. The key observation is that these log-likelihoods are, up to a constant, Bregman Divergences, for which the value of the mean parameter minimizing the average divergence of a dataset is the mean of that dataset (See Proposition 1 and Section 4.3 of Banerjee *et al.*^29^).

## Data

### Neurons

Data for 1.3 million mouse neurons were downloaded from 10X Genomics (22). We selected cells with at least 20,000 UMIs and subset to the 2000 most variable genes yielding a 4581 × 2000 count matrix. This matrix was subsampled to 3000 UMIs per cell to simulate a low-depth experiment and then processed as described in the Models section below.

### Myeloid Bone Marrow

Data for 3072 mouse cells from Paul, Arkin, & Giladi *et al.* (30) were downloaded from GEO (accession: GSE72857) based on the “Unsorted myeloid” label in the experimental design file. The data were filtered to cells with at least 1000 UMIs and to genes present in at least 10 cells, yielding a 2417 × 10783 count matrix. Hierarchical clustering in Fig. 2a was performed using the Scipy package^31^ with average linkage and Euclidean distance.

### EMT

Data for 7523 cells exhibiting the epithelial-to-mesenchymal transition from van Dijk *et al.* (19) were downloaded from the MAGIC Github repository^§^. The data were filtered to cells with at least 1000 UMIs and to genes in at least 10 cells, yielding a 7523 × 18259 count matrix. In Figure 3, the data is row-normalized to counts-per-N, where N is the median number of gene counts per cell.

### Simulated Dataset

We create a simulated dataset of 8 classes of 512 cells each, made up of 512 gene features. First we generate a matrix **P** of expression “programs” to transform points from an 8-dimensional latent space into 512-dimensional gene expression space. The class matrix **W** is defined as a random weighting over programs for each class. Multiplying these matrices into gene space yields a ground truth expression matrix **C** (in log space) that reflects the structure of the latent space and the class relationships. The expression *e*_*ij*_ for cell *i* from class *j* is generated by adding normally distributed noise to the mean expression of class *j*.

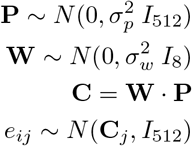

With 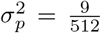 and 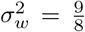 in these simulations. **P** is 8 × 512, **W** is 8 × 8, **C** is 8 × 512, and the expression matrix **E** is 4096 × 512.

UMI counts are sampled from this expression matrix using a variable library size, yielding a count matrix with 38% non-zero values. This level of sparsity is comparable to that found after restricting a single-cell dataset to deeply sequenced cells and relatively highly expressed genes. The count matrix is partitioned into two independent samples, and ground truth accuracy is assessed by comparison to the expected mean counts for each cell based on its library size and expression levels. The code used for generating simulated scRNA-seq data is available in www.github.com/czbiohub/simscity.

## Models

### PCA

The rank *k* matrix which is closest in the sense of mean-square error to a given matrix *X* is given by projecting *X* onto the span of the first *k* principal components. Concretely, if we write *X* = *U*Σ*V* for the singular value decomposition, then the denoised matrix is defined by *f*_*k*_(*X*):= *U*_*k*_Σ_*k*_*V*_*k*_, using the first *k* columns of *U*, the first *k* diagonal elements (singular values) of Σ, and the first *k* rows of *V*. When performing PCA we square-root normalized the UMI count matrix.

### Diffusion

We use a simple version of a diffusion model, which effectively averages similar expression vectors together. Concretely, we form a symmetrized 15-nearest neighbor graph *G* on the set of cells, where the distances used to determine neighbors are Euclidean distances in the 30-PC projection of the square-root and row-normalized gene expression matrix *X*. Let *W* be the transition matrix of a lazy random walk (go to a random neighbor with probability 0.5 and stay put with probability 0.5). For a given normalization function *η* and diffusion time *t*, the output of the denoiser is *f*_*t*_(*X*) = *W*^*t*^*η*(*X*). For mean-square error the count matrices were square-root normalized before diffusion, while for Poisson loss the raw counts were used.

This is a simplified implementation of the diffusion idea used in MAGIC, which includes adaptive neighborhood sizes, reweighted edges, and performs the diffusion in PC-space.

### Autoencoder

We use a simple autoencoder architecture where the encoder and decoder each have a single hidden layer, and are connected by a bottleneck layer which forces the autoencoder to compress the data. For the *Neuron* and *Myeloid* datasets the encoder and decoder hidden layers contained 512 nodes, while for the relatively simple simulated data they contained 128 nodes. All layers were fully-connected and used ReLU activation. For mean-square error the count matrices were square-root normalized, while for Poisson loss the input was log normalized (specifically *log*_*e*_(*x* + 1)).

Each network was trained using stochastic gradient descent with aggregated momentum^32^, with multiple cycles of cosine annealing until validation loss stopped improving. For complete details and code see https://www.github.com/czbiohub/molecular-cross-validation.

We note that training a deep-learning model can involve tuning many hyperparameters beyond the bottleneck size and network architecture, and while the results shown here illustrate the utility of molecular cross-validation for model selection, other architecture choices may obscure the relationship between model complexity and the self-supervised loss^33^. For example, in this work the autoencoder did not outperform PCA in spite of being a strictly more expressive model, highlighting that the training process is an integral part of a deep learning model.

---

**Figure S1:**
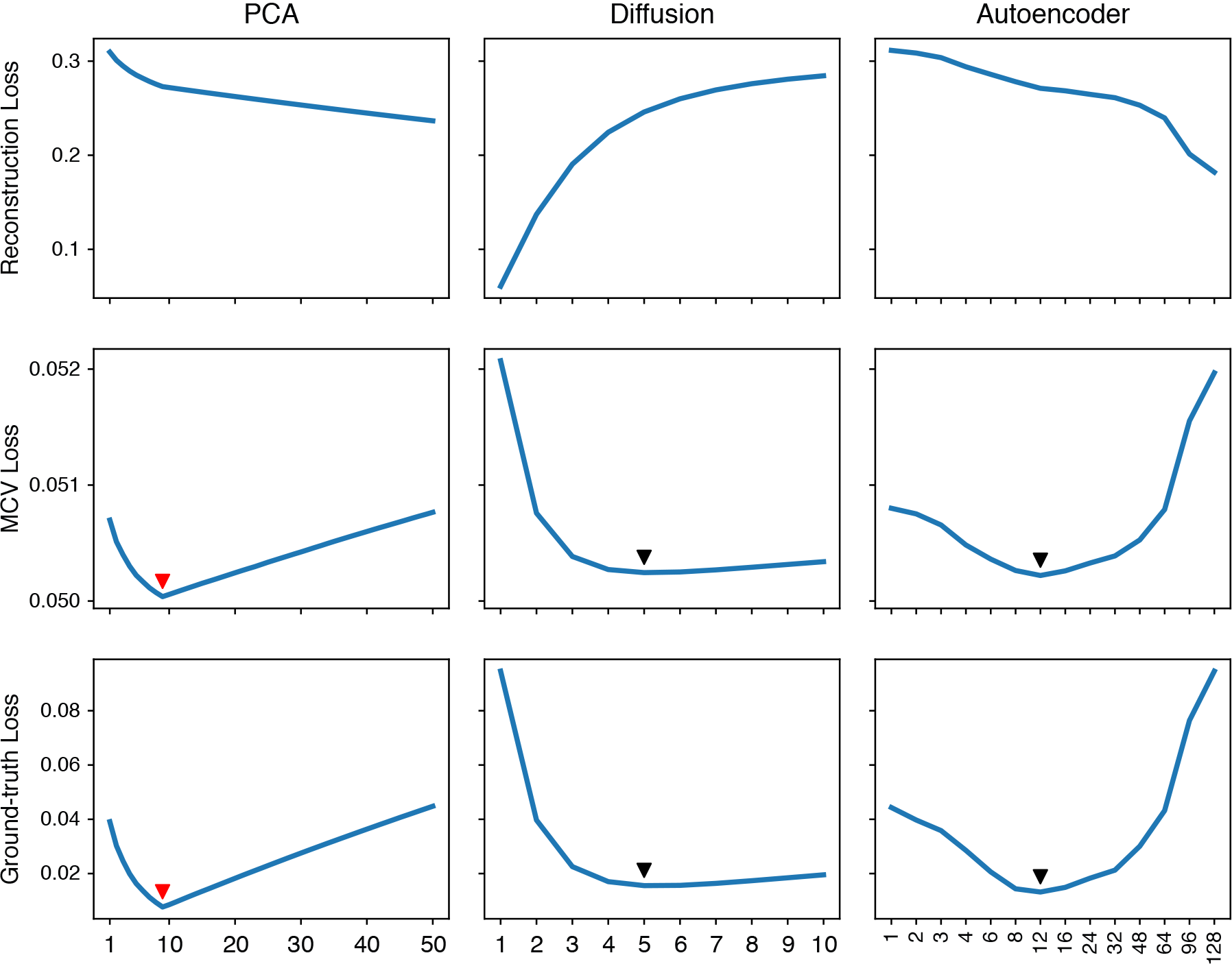
Performance of three denoising methods across a range of model parameters on simulated data. Arrows denote the minima of the MCV and ground-truth loss curves, which coincide. Red arrows denote the global minima among all three methods. For this dataset, the best model is PCA with 9 principal components.

**Figure S2:**
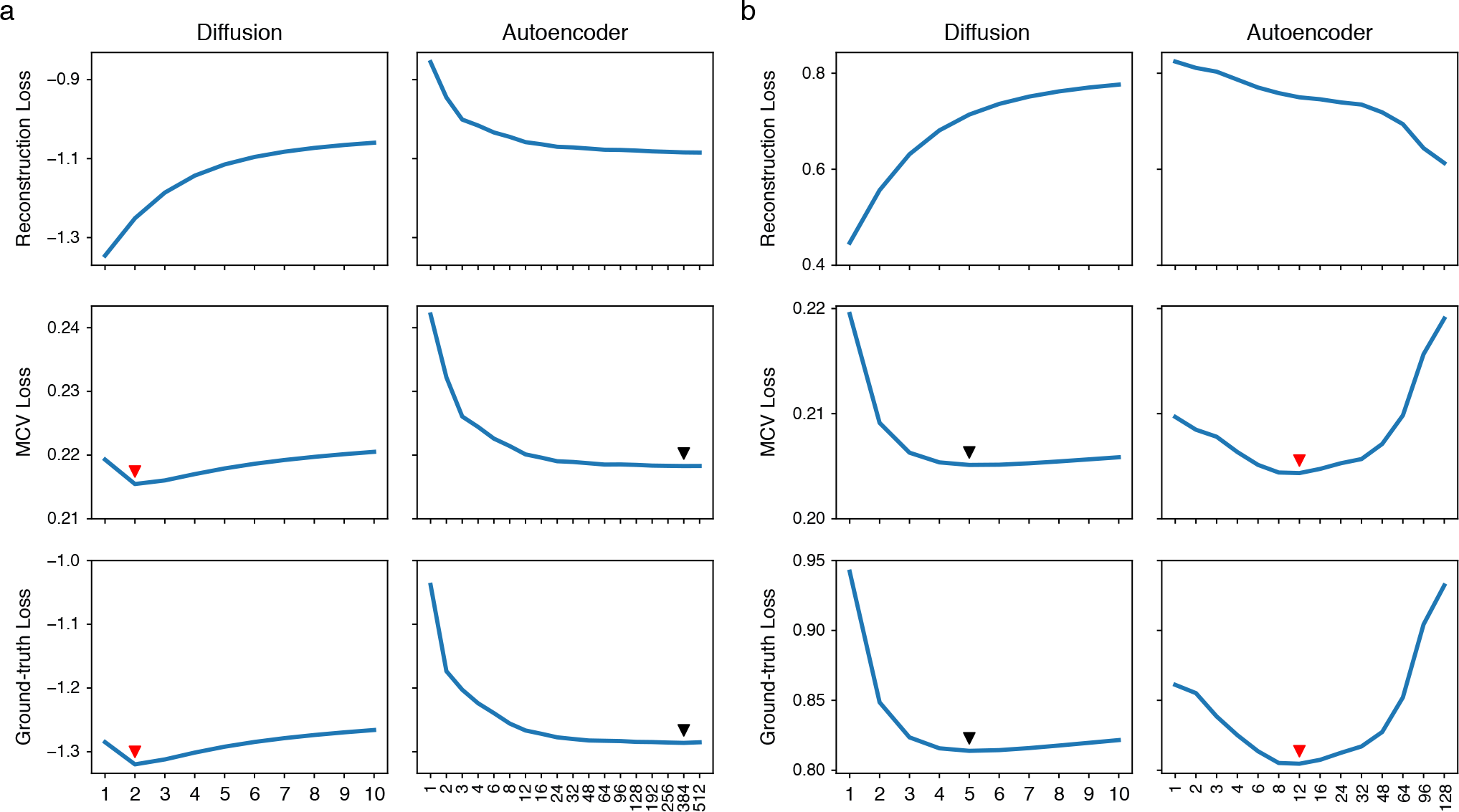
Performance of two denoising methods across a range of model parameters, evaluated using a Poisson loss. Arrows denote the minima of the MCV and ground-truth loss curves, which coincide. Red arrows denote the global minima. **(a)** On the *Neuron* dataset, diffusion with *t* = 2 performs the best. **(b)** On the simulated dataset, the autoencoder with bottleneck width 12 performs the best.

## Footnotes

∗ PCA can be viewed as a linear autoencoder with one hidden layer, the bottleneck, whose width is the number of principal components.

† Asymptotically, their shapes will be identical; with finite data they will merely be very close. See methods for details.

‡ For linguistic convenience we will assume that the counts are indexed by genes, but the same arguments apply if (pseudo)alignments to a transcriptome are used instead.

§ www.github.com/KrishnaswamyLab/MAGIC/tree/master/data

